# Theta oscillations in the prefrontal-hippocampal circuit do not couple to respiration-related oscillations

**DOI:** 10.1101/2021.12.22.473834

**Authors:** Sunandha Srikanth, Dylan Le, Yudi Hu, Jill K. Leutgeb, Stefan Leutgeb

## Abstract

Oscillatory activity is thought to coordinate neural computations across brain regions, and theta oscillations are critical for learning and memory. Because the frequency of respiratory-related oscillations (RROs) in rodents can overlap with the frequency of theta in the prefrontal cortex (PFC) and the hippocampus, we asked whether odor-cued working memory may be supported by coupling between these two oscillations. We first confirmed that RROs are propagated to the hippocampus and PFC and that RRO frequency overlaps with canonical theta frequency. However, we found low coherence between RROs and local theta oscillations in the hippocampus-PFC network when the two types of oscillations overlapped in frequency. This effect was observed during all behavioral phases including during movement and while odors were actively sampled when stationary. Despite the similarity in frequency, RROs and theta oscillations therefore appear to be limited to supporting computation in distinct networks, which suggests that sustained long-range coordination between oscillation patterns that depend on separate pacemakers is not necessary to support at least one type of working memory.

## INTRODUCTION

Brain oscillations are thought to coordinate neural computations across cortical and sub-cortical brain regions by synchronizing network activity (Buzsáki & Draguhn, 2004). In brain circuits that support memory function, such coordination is most prominent in the theta frequency range (6-12 Hz). Theta oscillations are not only prominent in the hippocampus, but matching oscillations can also be observed in directly and indirectly connected brain regions (Backus et al., 2016; Buzsáki, 2002; Colgin, 2011). For example, local field potentials (LFP) are highly coherent between hippocampus and medial prefrontal cortex (mPFC), and neuronal firing patterns of many prefrontal neurons are phase-locked to the hippocampal theta rhythm (Hyman et al., 2005; Jones & Wilson, 2005; Siapas et al., 2005; Zielinski et al., 2019). Since oscillations within a network correspond to cyclic fluctuations in excitability, such synchronized oscillations across brain regions allow for windows of peak excitability that enable efficient communication between the brain regions (Fries, 2005). Accordingly, the accuracy of spatial coding in hippocampus and mPFC has been reported to be coupled on a cycle-by-cycle basis (Zielinski et al., 2019). Furthermore, prefrontal-hippocampal oscillatory strength correlates with performance in spatial working memory tasks in rodents (Benchenane et al., 2010; Jones & Wilson, 2005; Zielinski et al., 2019), which suggests that oscillatory coupling supports memory function and raises the question whether an even broader network is dynamically synchronized during task performance.

Along with the canonical theta oscillations that are most prominent in the hippocampus, oscillations that encompass an overlapping frequency range (3-12 Hz) and are related to the respiration rhythm have also been described (Lockmann et al., 2016; Nguyen Chi et al., 2016; Rojas-Líbano et al., 2014; Yanovsky et al., 2014). Respiration is paced by brainstem breathing centers (Feldman et al., 2013), and the nasal airflow that is generated by breathing then activates olfactory sensory neurons in the nasal epithelium during each breathing cycle (Wu et al., 2017). This mechanism entrains local oscillatory activity in the olfactory bulb (OB), and the respiratory rhythm and OB oscillations are thus tightly coupled. In particular, a causal role of nasal airflow for olfactory oscillations has been established by the finding that the entrainment of OB network activity is diminished when nasal airflow is restricted by means of naris occlusion or tracheal breathing (Onoda & Mori, 1980; Phillips et al., 2012).

Respiration-entrained activity of OB neurons is transmitted to olfactory-associated cortical areas such as the piriform cortex (Fontanini et al., 2003) and the barrel cortex (Ito et al., 2014), but also to more indirectly connected subcortical and cortical regions across the brain, including the medial prefrontal cortex (mPFC) (Biskamp et al., 2017) and the hippocampus (Lockmann et al., 2016; Nguyen Chi et al., 2016; Tort et al., 2018; Yanovsky et al., 2014). Throughout these brain regions, respiration-related oscillations (RROs) can be distinguished from other types of oscillations by confirming the coupling to either the respiration rhythm and/or olfactory bulb oscillations. Consistent with the definition of RROs as respiration or OB-oscillation related, these oscillatory patterns in the mPFC, barrel cortex and the hippocampus are disrupted when manipulating signals from the OB through bulbectomies and olfactory epithelial ablation or when disturbing nasal airflow through tracheotomies and naris occlusions (Biskamp et al., 2017; Ito et al., 2014; Moberly et al., 2018; Nguyen Chi et al., 2016; Yanovsky et al., 2014).

Because the overlap in frequency between RROs and canonical theta can be confounding for separately analyzing these oscillation patterns, characterization of RROs has mostly focused on periods when RROs differ in frequency from theta oscillations during running, immobility and anesthesia (Nguyen Chi et al., 2016; Yanovsky et al., 2014). In these analyses, oscillations at the respiratory frequency have a different depth profile than theta oscillations in the hippocampus, which supports the notion that RROs are separate oscillations that co-occur with theta oscillations in the hippocampus. In contrast, there is also evidence that olfactory oscillations and hippocampal theta oscillations become coherent during periods of sniffing in odor learning and discrimination tasks (Kay, 2005; Macrides et al., 1982). These latter studies suggest that the coherence between hippocampal and olfactory networks mediates sensorimotor integration in the hippocampus. A possible source for the conflicting reports on the coupling of respiratory oscillations and hippocampal oscillations is that these reports have not considered the existence of two types of theta oscillations in the hippocampus – movement-related theta oscillations and sensory-evoked theta oscillations (Kramis et al., 1975; Vanderwolf, 1969). We therefore investigated whether coupling between RROs and theta oscillation may differ depending on the behavioral state during which theta is generated. Furthermore, we reasoned that coupling between the respiratory rhythm and hippocampal canonical theta may be required when olfactory cues are relevant for memory performance and performed recordings in an odor-cued hippocampus-dependent working memory task. To be able to identify RROs throughout the behavior, we recorded OB oscillations simultaneously with hippocampal oscillations. Furthermore, we also simultaneously recorded from mPFC to examine whether the convergence of RROs and hippocampus-coupled theta in mPFC would allow for dynamic coupling between these two types of oscillations, which could in turn serve as a conduit for coordinating memory and sensory processing in the prefrontal-hippocampal circuit.

## RESULTS

To investigate the coupling between the respiration-coupled oscillations in the OB and theta oscillations in the mPFC and hippocampus, we simultaneously recorded LFP signals across these brain regions. Because ventral hippocampus (vHC) is connected more strongly to mPFC than dorsal hippocampus (dHC) (Hoover & Vertes, 2007), we placed separate recording electrodes in the dHC and the vHC. Within mPFC, we focused on the prelimbic, infralimbic, and anterior cingulate areas because of their direct and indirect connections with hippocampus. RROs as well as theta oscillations have been detected in all of these regions in previous studies (Tort et al., 2018).

To be able to examine oscillations across a range of behavioral states, we trained mice in an odor-cued working memory task (**Figure 1A**) in which we could observe movement-related hippocampal theta while mice ran between an odor port and reward locations and sensory-related theta when mice sampled odors while they remained stationary at the odor port. Briefly, mice (*n* = 8) were trained to run on a figure-8 maze in which an odor port was placed at one end of the stem arm where one of two odors (isoamyl acetate or ethyl acetate) was delivered in a pseudorandom fashion. Mice were trained to sample the odor by poking and holding their nose in the odor port for at least one second. They then had to retain information about the odor identity while running to the opposite end of the stem arm to make their choice to turn left or right. Correct choices were rewarded with a single chocolate sprinkle. As expected, initial performance was at chance level (*n* = 8 mice, Z = 0.49, p = 0.31, one-tailed Wilcoxon signed-rank test). Using a criterion of 65% correct during at least 2 of 3 consecutive days, mice learned the task within 15 ± 5 days. Accordingly, performance during the last three days of testing was better than during the first three days (median: 69.5% vs 50.2% correct, *n* = 8 mice, Z = 2.45, p = 0.007, one-tailed Wilcoxon signed-rank test; **Figure 1B**). All analyses of electrophysiological data were performed on data from the last three testing days for each animal. Recording sites in the OB, mPFC, dHC and vHC were confirmed in histological material (**Figure 1C**). Since histological confirmation of electrode locations was not successful in one animal, we included only 7 of 8 animals for all LFP analysis.

**Figure 1.**
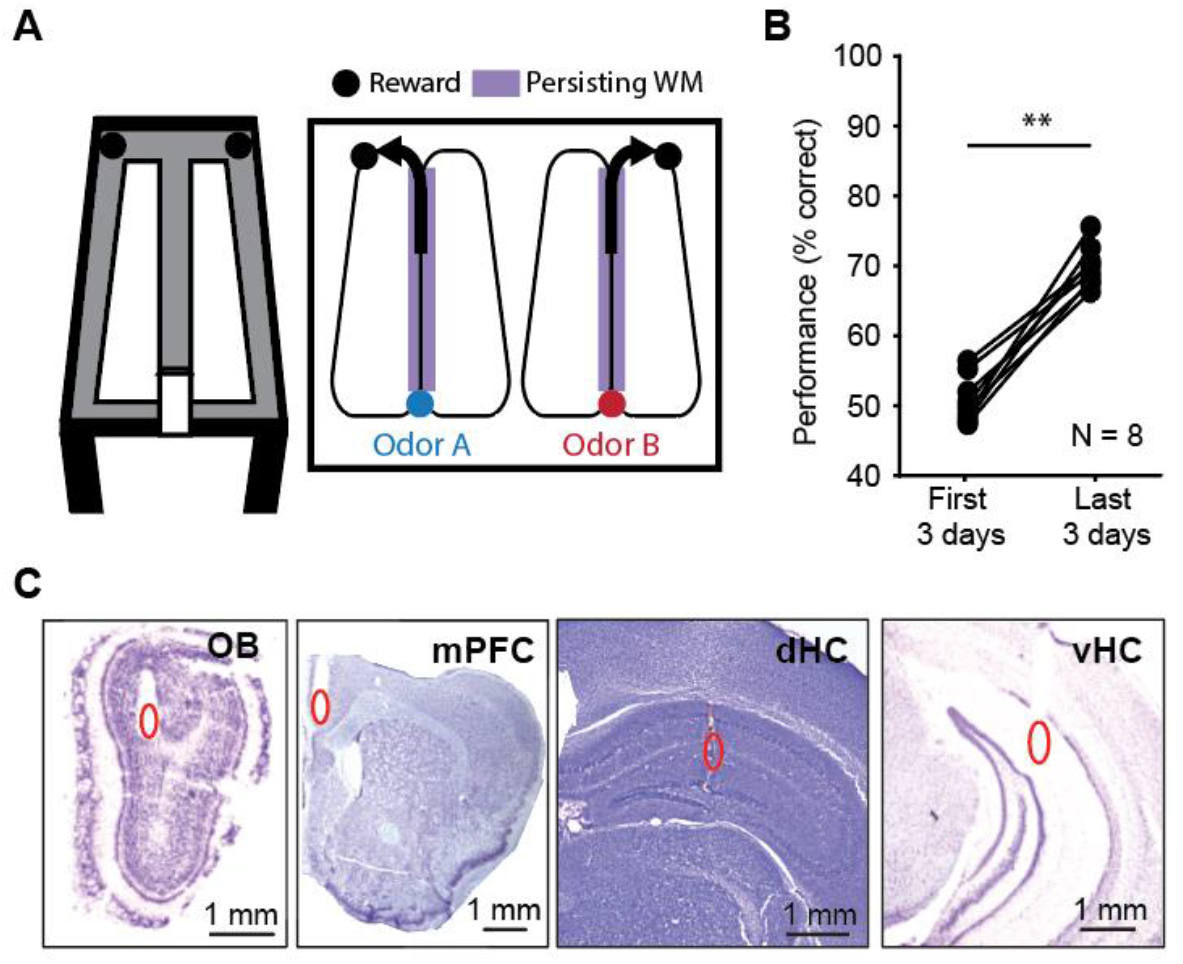
Mice performed an odor-cued working memory task with high accuracy. **A**. Schematic of the odor-cued working memory task. Mice were trained to sniff one of two pseudo-randomly delivered odors at an odor port at the bottom of the stem arm and make a turn at the top of the stem arm based on the odor they sampled. The relation between odor identity and turn direction remained consistent for each mouse. A food reward was provided at the reward zones for correct choices, and mice returned to the odor port on the side arms. **B**. Performance increased between the first three days and the last three days of behavioral testing (*n* = 8 mice, Z = 2.45, p = 0.007, one-tailed Wilcoxon signed-rank test) **C**. Example recording electrode locations in the olfactory bulb (OB), medial prefrontal cortex (mPFC), dorsal hippocampus (dHC) and ventral hippocampus (vHC). Recording locations are highlighted (red ovals) in cresyl-violet stained brain slices.

### Predominant OB frequencies ranged from 3-12 Hz in all task phases

The task was parsed into four task phases with distinct behavioral patterns – return arm where animals ran without a memory load, odor sampling when animals actively sampled an odor, stem arm where animals ran after odor sampling and before making a choice, and reward arm where animals were rewarded for correct performance. Time periods when animals transitioned between these phases were not considered. During the odor sampling period, the animals poked their noses into the odor port and sampled the odor while holding the nose in the odor port and were thus stationary. For meaningful comparisons with the stationary odor sampling periods, another period with minor movements was selected by restricting analyses on the reward arm to periods with low velocity (less than 5 cm/s). Conversely, we confirmed that running speeds on the return arms and on the stem were high and matched (**Figure 2A**), which allowed us to compare two task phases with corresponding movement patterns.

**Figure 2.**
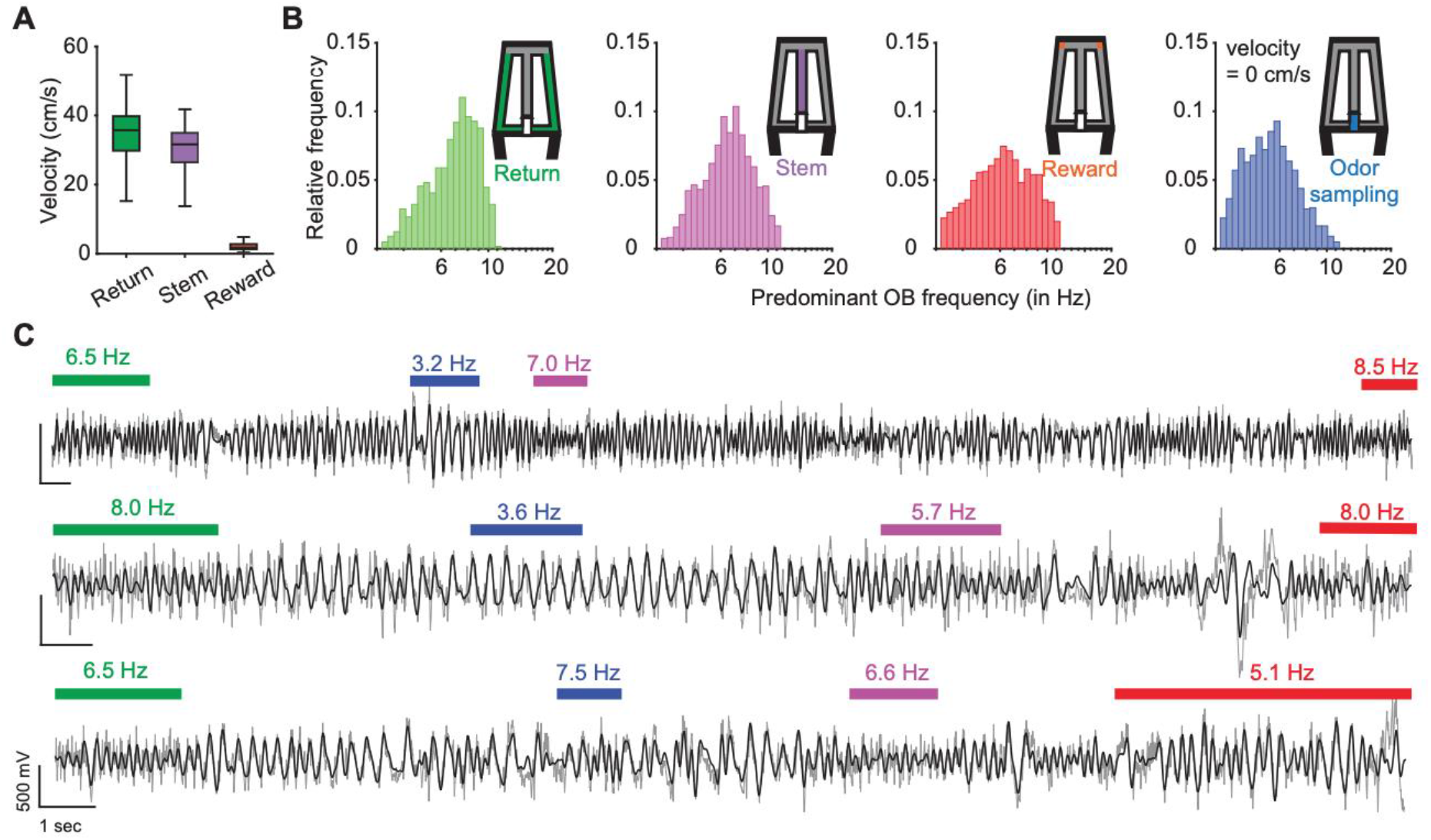
OB frequencies within each task phase ranged from 3-12 Hz. **A**. Velocity of the mice (*n* = 7 mice, 1207 trials) in each maze zone. Velocity in the odor port is not shown and was near zero while animals had to hold their nose in the port during odor sampling. In the box plots, the center line shows the median, and the bottom and top edges of the box represent the 25th and 75th percentiles, respectively. The whiskers indicate the most extreme data points. **B**. The predominant OB frequency in each trial was calculated and the distribution of predominant OB frequencies across trials is plotted for each task phase. **C**. Example OB LFP traces (grey: raw traces, black: 3-12 Hz filtered traces). Each line is a trial, and colored bars indicate time periods when animals were in the respective task phase (green: return arm, blue: odor sampling, purple: stem arm and red: reward zone). Numbers on top of bars indicate the predominant OB frequency. Transition phases are without bars and were not analyzed.

Although the velocity profiles ranged from immobility in the odor port to high running speeds on the return arms (**Figure 2A**), the frequency distributions of OB oscillations across the four task phases showed only minor differences (*n* = 1207 trials, median±iqr in return: 7.02±2.77 Hz; stem: 6.58±2.60 Hz; reward: 6.16±3.26 Hz; odor sampling: 5.41±2.42 Hz), and the entire frequency range from 3 Hz to 12 Hz was observed across trials for each of the phases (**Figure 2B**). Moreover, within a trial, predominant OB frequencies varied across task phases (**Figure 2C**) with only weak correlations among them (Spearman correlation coefficients <0.5) (**Figure 2 – figure supplement 1**). OB oscillations in our analyzed frequency range have been firmly established as being generated by respiration (Jessberger et al., 2016; Phillips et al., 2012; Rojas-Líbano et al., 2014), and we thus refer to these oscillations as respiratory-related oscillations (RROs).

### RROs and canonical theta differed in their frequency distributions

Because our task design included phases with running and immobility, it allowed us to assess the coordination of RROs with theta across these task phases. As expected for movement-related theta, high amplitude theta oscillations were observed during periods of running on the stem and return arms (**Figure 3A** and **Figure 3 – figure supplement 1A**). Hippocampal theta oscillations were also observed during odor sampling, and because mice were stationary while holding the nose in the odor port, theta oscillations during this task phase can be considered sensory evoked (**Figure 3A** and **Figure 3 – figure supplement 1B**). Sensory-evoked theta oscillations were lower in amplitude than movement-related theta oscillations (**Figure 3A**) but could nonetheless be clearly detected in the dHC as distinct from RROs based on their frequency distribution (**Figure 3B** and **Figure 3 – figure supplement 1C**). The predominant frequencies of the hippocampal oscillations during odor sampling were as narrowly distributed as during movement, while RROs varied more widely in frequency than the hippocampal oscillations (**Figure 3B** and **Figure 3 – figure supplement 1C** and **D**). Hippocampal theta oscillations were also detected on reward arms, but this period included bouts with low movement velocity (<5 cm/s) such that the type of theta cannot be as clearly classified as during other phases of behavior. Irrespective of behavior phase, the distribution of predominant frequencies in OB therefore included the entire 3-12 Hz range while canonical dHC theta was mostly concentrated in the 7-11 Hz range (**Figure 3 - figure supplement 1C**, **Figure 3B**).

**Figure 3.**
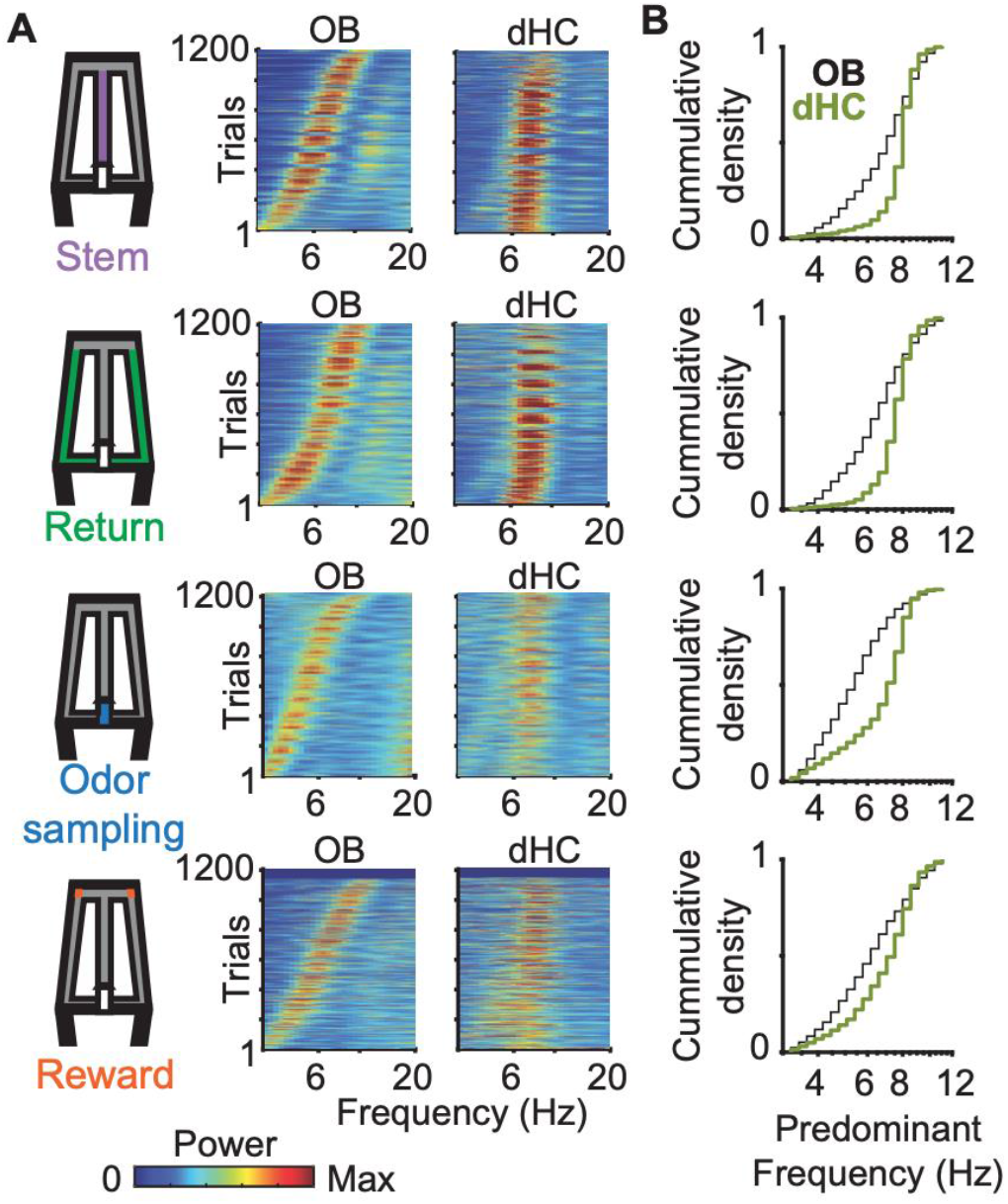
Across trials, predominant OB oscillation frequencies and dHC oscillation frequencies varied so that they were either overlapping or non-overlapping. **A.** Power spectra of OB and dHC oscillations are shown as color-coded plots, with each line corresponding to a trial. Trials are ordered by the OB peak oscillation frequency. B. Cumulative density functions of the predominant OB frequencies (black) and dHC frequencies (green) for the four task phases. The data for OB frequencies is replotted from Figure 2A for comparisons with dHC frequencies. Predominant dHC frequencies were concentrated in the range of 7-11 Hz, while OB frequencies spanned the entire range of 3-12 Hz during all the task phases. Frequency distributions differed between brain regions in all task phases (*n* = 1207 trials, Return: p = 3.4e-56, KS = 0.32; Stem: p = 1.8e-70, KS = 0.36; Odor sampling: p = 7.1e-88, KS = 0.41; Reward: p = 4.7e-21, KS = 0.20).

The wider frequency range for OB compared to canonical theta oscillations implied that there were trials in which the predominant OB frequency and the canonical dHC theta frequency either differed or overlapped. We therefore grouped trials into two categories – trials with overlapping dHC theta and OB frequency (≤1 Hz apart) and trials with non-overlapping dHC theta and OB frequency (>1 Hz apart) (**Figure 4**). Grouping of trials as overlapping or non-overlapping in frequency was done independently for each task phase within a trial. For example, a trial could be grouped as overlapping for analysis on the stem arm and as non-overlapping for analysis on the return arm.

**Figure 4.**
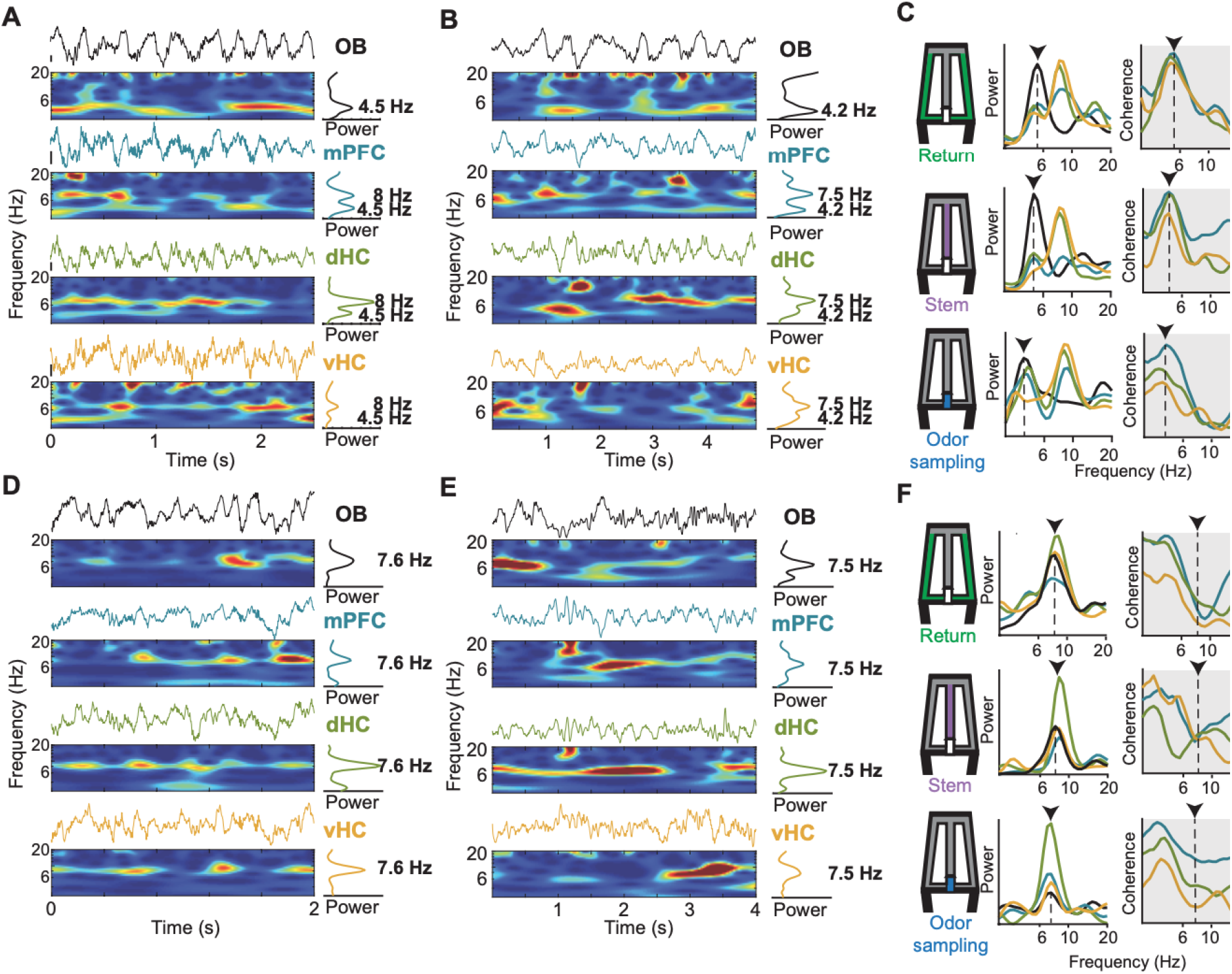
RROs were observed in the mPFC-dHC-vHC network in parallel with movement-related or sensory-evoked theta oscillations. **A**. Example raw traces and corresponding time-frequency spectrograms of simultaneously recorded LFP from the OB, mPFC, dHC and vHC are shown for a period when the animal was running on the stem arm of the maze. In the example, theta frequency in the mPFC-dHC-vHC regions was non-overlapping with the predominant OB frequency. **B**. Arranged as in A but for an example period when the mouse was stationary while actively sampling odor at the port. As in A, theta and OB frequencies were non-overlapping. **C**. Time averaged power spectra (left) and coherence spectra (right) are shown for three example periods within a trial and maze zone (return arm, stem arm and odor sampling period) when OB and canonical theta frequencies were non-overlapping. Dotted lines and arrows indicate the frequency of the predominant OB oscillation in the respective trials. OB-mPFC, OB-dHC and OB-vHC coherence is higher at the frequency matching the predominant OB frequency compared to the theta frequency. **D**, **E**. Arranged as in A and B, respectively, but for example periods when OB and canonical theta oscillations overlapped in frequency. **F**. Arranged as in C but for example periods (return arm, stem arm and odor sampling period) with overlapping OB and canonical theta frequencies. Despite the similar peak frequencies of both types of oscillations, OB-mPFC, OB-dHC and OB-vHC coherence was low at the frequency matching the predominant OB frequency. Accordingly, canonical theta oscillations in the mPFC-dHC-vHC network did not couple to respiration-entrained oscillations.

### RROs were observed in the prefrontal-hippocampal network in parallel with either movement-related or sensory-evoked theta oscillations

We first investigated the coordination between theta oscillations and RROs in trials with non-overlapping frequencies. Theta oscillations are well defined as movement-related during periods of running on the stem and running on return arms and as sensory-evoked during odor sampling. The analysis therefore focused on these periods and excluded reward. During running on the stem and return arms, LFP signals at the prefrontal and hippocampal recording sites showed two detectable peaks in the 3-12 Hz range – one at a frequency of ∼8 Hz and another matching the predominant OB frequency (**Figure 4A** and **C**). While being stationary during odor sampling periods, LFP in the prefrontal-hippocampal network showed the same two peaks (**Figure 4B** and **C**). Given that clear peaks that matched the OB frequency could be detected in mPFC, dHC, and vHC, we asked whether the peak at the OB frequency indicated coupled oscillations across all brain regions. We measured coupling by computing the coherence spectra between OB oscillations and prefrontal as well as hippocampal oscillations. Consistent with coupled RROs across brain regions, the maximum of the coherence spectra between the OB and each of the other recording sites was at frequencies matching the predominant OB frequency. These results indicate that broadly transmitted RROs were clearly detectable across brain regions and even at dorsal hippocampal recording sides where canonical theta oscillations showed a substantially higher amplitude than RROs (**Figure 4C**).

Since most trials with nonoverlapping OB and dHC frequencies were those in which the predominant OB frequency was below 6 Hz, we asked whether a comparison of non-overlap trials with OB frequency ≥6 Hz would yield similar results as for those below 6 Hz. We found that coherence was significantly lower in non-overlap trials with OB frequency ≥6 Hz compared to trials with OB frequency <6Hz for all combinations of OB oscillations and cortical regions and for all maze segments (return arm: *n* = 365 and 378 trials with OB frequency <6 Hz and ≥6 Hz, OB-mPFC: Z = -14.70, p = 6.3e-49; OB-dHC: Z = -15.91, p = 4.9e-57; OB-vHC: Z = -13.16, p = 1.3e-39, Wilcoxon signed-rank test; stem: *n* = 447 and 305 trials, OB-mPFC: Z = -8.00, p = 1.2e-15; OB-dHC: Z = -10.31, p = 6.2e-25; OB-vHC: Z = -5.99, p = 2.0e-9, Wilcoxon signed-rank test; odor sampling period: *n* = 776 and 212 trials, OB-mPFC: Z = -11.57, p = 5.5e-31; OB-dHC: Z = -10.75, p = 5.4e-27; OB-vHC: Z = -7.22, p = 5.1e-13, Wilcoxon signed-rank test) (**Figure 4 – figure supplement 1**). OB oscillations at high frequencies – when differing from the frequency of the predominant hippocampal and prefrontal theta oscillations – therefore do not appear to couple across brain regions.

### Movement-related and sensory-evoked theta oscillations did not couple to RROs

While it is expected that RROs and canonical theta (either movement-related or sensory-evoked) are unlikely to couple when their frequencies are non-overlapping, we considered it plausible that RROs showed coupling with either type of theta oscillation when the frequencies were matching. By selecting trials in which predominant OB frequencies and hippocampal frequencies overlapped (**Figure 4D** and **E**), we were able to ask whether RROs couple to canonical theta oscillations in mPFC, dHC or vHC regions. If theta oscillations in the prefrontal-hippocampal network were to couple to RROs in this trial type, we would expect a high coherence. In contrast, we found a significant decrease in coherence in trials with overlapping frequencies compared to trials with non-overlapping frequencies for all combinations of OB oscillations and cortical regions and for all maze segments (return arm: *n* = 743 non-overlap and 464 overlap trials, OB-mPFC: Z = -10.79, p = 3.7e-27; OB-dHC: Z = -9.09, p = 9.6e-20; OB-vHC: Z = -8.54, p = 1.3e-17, Wilcoxon signed-rank test; stem: *n* = 752 and 455 trials, OB-mPFC: Z = -7.67, p = 1.7e-14; OB-dHC: Z = -4.88, p = 1.0e-6; OB-vHC: Z = -4.82, p = 1.4e-6, Wilcoxon signed-rank test; odor sampling period: *n* = 956 and 251 trials, OB-mPFC: Z = -10.09, p = 6.4e-24; OB-dHC: Z = -7.34, p = 2.1e-13; OB-vHC: Z = -6.61, p = 3.9e-11, Wilcoxon signed-rank test; **Figure 4F** and **5**). The decrease in coherence in trials with overlapping compared to non-overlapping frequencies could not be explained by a difference in power of these oscillations between trial types (return arm: *n* = 743 non-overlap and 464 overlap trials, mPFC: Z = 10.73, p = 1.00; dHC: Z = 19.15, p = 1.00; vHC: Z = 13.59, p = 1.00, one-tailed Wilcoxon signed-rank test; stem: *n* = 752 and 455 trials, mPFC: Z = 8.58, p = 1.00; dHC: Z = 21.51, p = 1.00; vHC: Z = 14.34, p = 1.00, one-tailed Wilcoxon signed-rank test; odor sampling period: *n* = 956 and 251 trials, mPFC: Z = 4.90, p = 1.00; dHC: Z = 14.34, p = 1.00; vHC: Z = 8.59, p = 1.00, one-tailed Wilcoxon signed-rank test). Furthermore, a comparison of running velocity in trials with overlapping and non-overlapping frequencies did not reveal any difference (return: *n* = 743 non-overlap and 464 overlap trials, Z = -1.44, p = .07, stem: *n* = 752 and 455 trials, Z = 7.21, p = 1.00, one-tailed Wilcoxon signed-rank test). Taken together, these results suggest that either type of canonical theta oscillation – movement-related or sensory-evoked – in cortical regions did not couple to the respiration-entrained oscillation in the OB during any of the behavioral periods.

**Figure 5.**
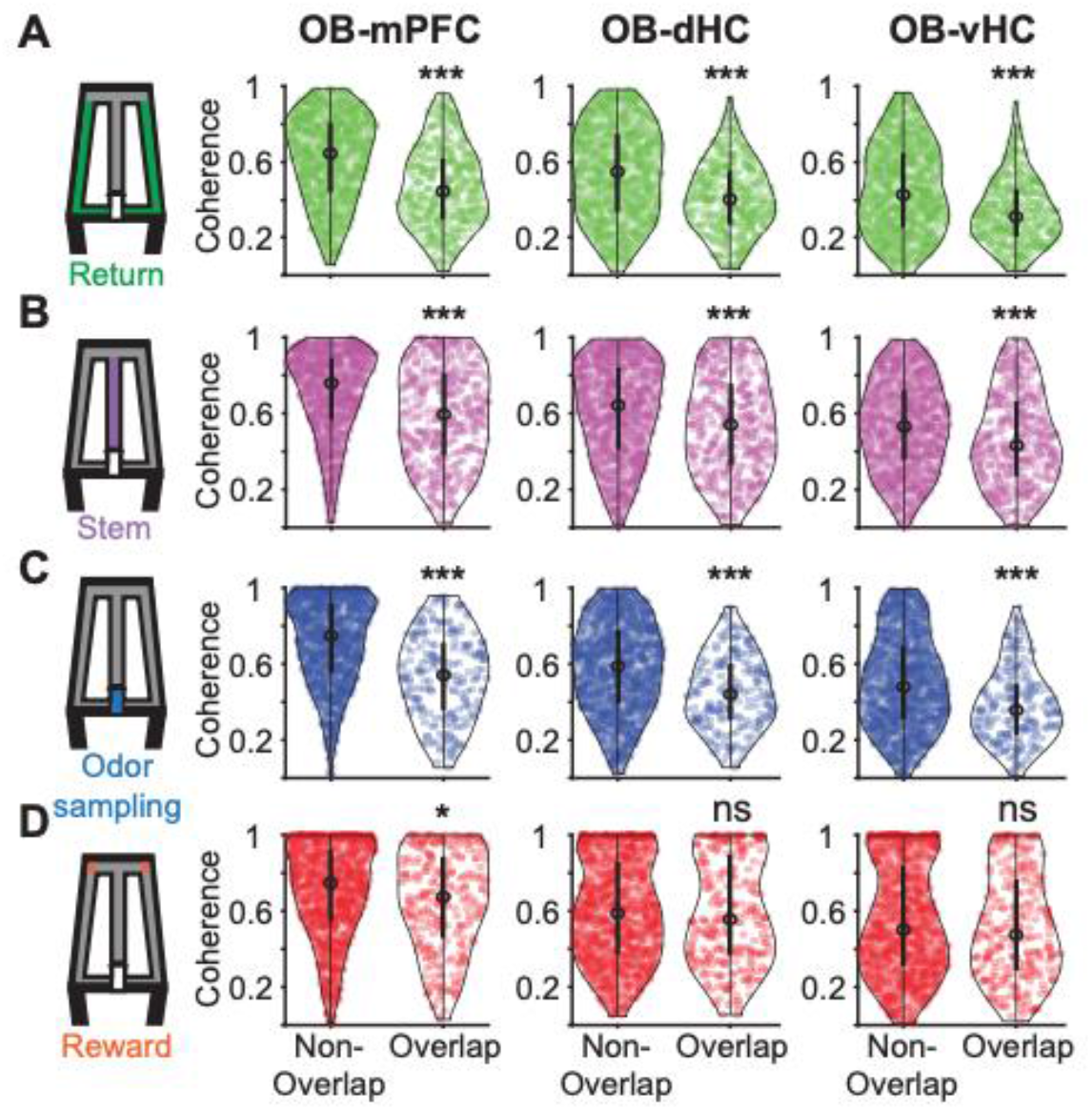
Respiration-entrained oscillations in the OB were not coupled to movement-related and sensory-evoked theta oscillations in the mPFC-dHC-vHC network. Violin plots of OB-mPFC, OB-dHC, and OB-vHC coherence in (**A**) the return arm, (**B**) the stem arm, (**C**) the odor sampling period and (**D**) the reward arm. Coherence was calculated at the RRO frequency in each trial, and trials with overlapping and non-overlapping theta and RRO frequencies were analyzed separately. A significant decrease in coherence was found in trials with overlapping RRO and theta frequencies compared to trials with non-overlapping frequencies for all prefrontal-hippocampal regions in the return arm (*n* = 743 non-overlap and 464 overlap trials, OB-mPFC: Z = -10.79, p = 3.7e-27; OB-dHC: Z = -9.09, p = 9.6e-20; OB-vHC: Z = -8.54, p = 1.3e-17, Wilcoxon signed-rank test), stem arm (*n* = 752 and 455 trials, OB-mPFC: Z = -7.67, p = 1.7e-14; OB-dHC: Z = -4.88, p = 1.0e-6; OB-vHC: Z = -4.82, p = 1.4e-6, Wilcoxon signed-rank test) and odor sampling period (*n* = 956 and 251 trials, OB-mPFC: Z = -10.09, p = 6.4e-24; OB-dHC: Z = -7.34, p = 2.1e-13; OB-vHC: Z = -6.61, p = 3.9e-11, Wilcoxon signed-rank test). In the reward zone, only OB-mPFC coherence (*n* = 917 and 290 trials, Z = -3.07, p = 0.0021, Wilcoxon signed-rank test) was decreased in trials with overlapping RRO and theta frequencies compared to trials with non-overlapping frequencies. In the violin plots, the center circle indicates the median, and the bottom and top of the thick black lines indicate the 25^th^ and 75^th^ percentile of the data respectively.

### Movement-related theta oscillations within prefrontal-hippocampal regions were more coherent in the stem compared to return arm

After establishing that RROs did not couple to either type of theta oscillations in any of the cortical regions, we explored whether we could nonetheless identify dynamic coordination of movement-related theta oscillations between cortical areas, which has previously been described in similar tasks. We analyzed the two task phases (stem and return arms) when movement-related theta oscillations were observed (**Figure 3 – figure supplement 1**). While the mPFC and dHC regions did not show an increase in movement-related theta power in the stem compared to the return arms (mPFC: *n* = 1207 trials, Z = 9.78, p = 1.00; dHC: *n* = 1207 trials, Z = 0.75, p = 0.22, one-tailed Wilcoxon signed-rank test), vHC theta power was higher in the stem compared to the return arms (vHC: *n* = 1207 trials, Z = -4.30, p = 5.82e-7, Wilcoxon signed-rank test; **Figure 6A** and **B**). Although there was a modest increase in movement-related theta power in only the vHC, coherence of movement-related theta oscillations between all three pairs of regions was higher in the stem compared to the return arms (mPFC-dHC: *n* = 1207 trials, Z = -14.34, p = 4.54e-47; mPFC-vHC: *n* = 1207 trials, Z = -11.49, p = 1.67e-29; dHC-vHC: *n* = 1207 trials, Z = -14.64, p = 3.93e-50, Wilcoxon signed-rank test; **Figure 6C** and **D**). This analysis does not only reproduce previous findings, but also confirms that our methods can, in principle, detect behavior-dependent increases in coherence.

**Figure 6.**
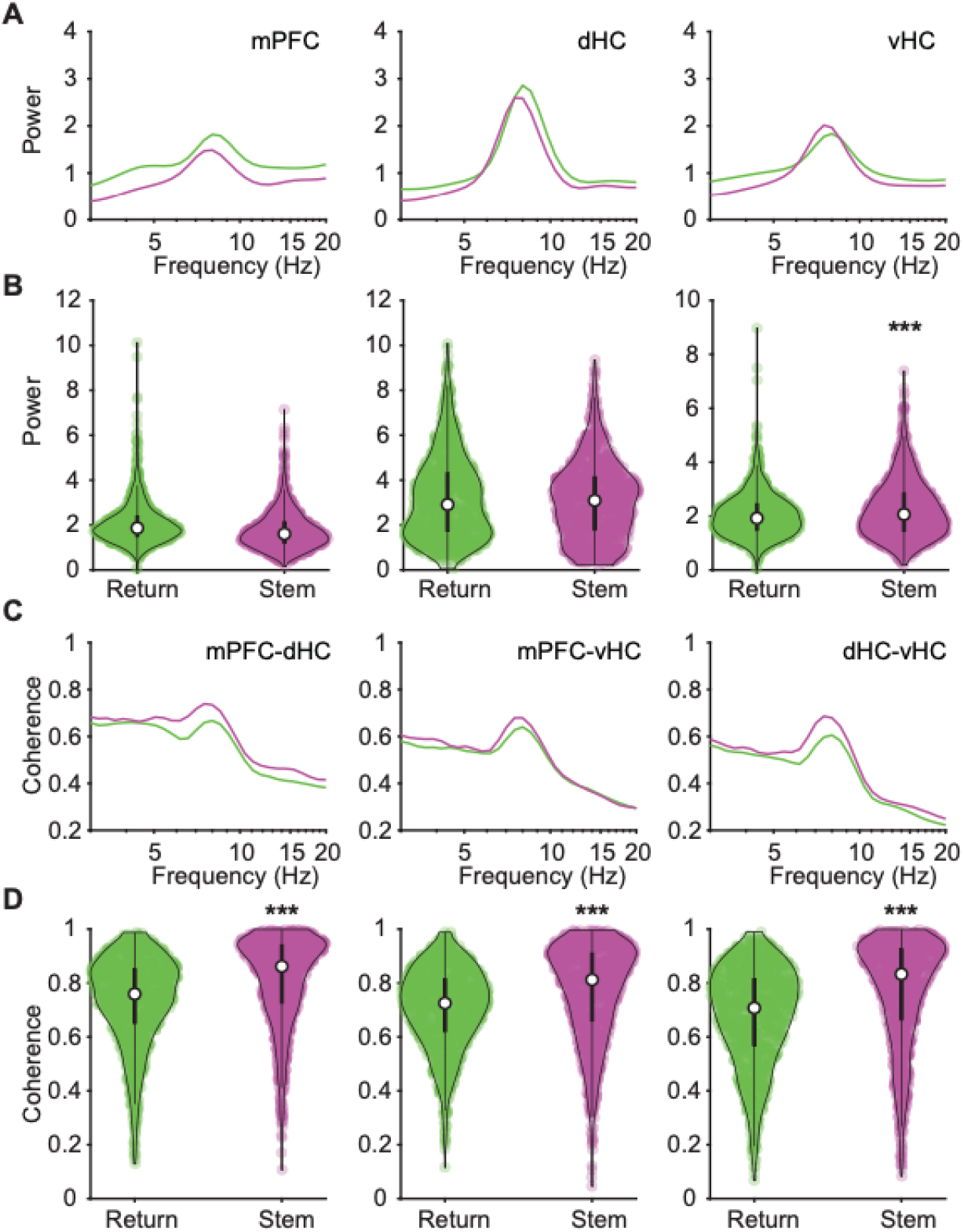
Movement-related theta oscillations in the prefrontal-hippocampal regions were more coherent in the stem arm compared to return arm. **A**. Trial-averaged power spectra of mPFC, dHC and vHC regions for periods on return arms (green) and on the stem arm (purple). **B**. Violin plots of peak movement-related theta power in the mPFC, dHC and vHC regions during periods on the stem and return arms across all trials. The mPFC and dHC regions did not show an increase in movement-related theta power in the stem arm compared to the return arm (*n* = 1207 trials, mPFC: Z = 9.78, p = 1.00; dHC: Z = 0.75, p = 0.22), but vHC theta power was higher in the stem arm compared to the return arm (vHC: Z = -4.30, p 5.82e-7, one-tailed Wilcoxon signed-rank test). **C**. Trial-averaged coherence spectra for pairs of regions (mPFC-dHC, mPFC-vHC and dHC-vHC). **D**. Violin plots of the coherence of movement-related theta oscillations. Coherence of all three pairs of regions was compared between the stem (purple) and return (green) arms. Coherence of movement-related theta oscillations was higher in the stem arm compared to the return arm (*n* = 1207 trials, mPFC-dHC: Z = -14.34, p = 4.54e-47; mPFC-vHC: Z = -11.49, p = 1.67e-29; dHC-vHC: Z = -14.64, p = 3.93e-50, Wilcoxon signed-rank test). In the violin plots, the center circle indicates the median, and the bottom and top of the thick black lines indicate the 25^th^ and 75^th^ percentile of the data, respectively.

### Coherence between prefrontal cortex and hippocampal regions was unrelated to odor-guided memory performance

Even though we examined odor-cued working memory in our task, the figure-8 maze is often used to assess spatial alternation behavior in rodents. While we did not train the mice to alternate in the odor-cued task, we observed spatial alternation on successive trials (i.e., right turn followed by a left turn or vice versa) in the beginning of behavioral training when the odor-cued choice behavior was at chance. We reasoned that mice’s propensity towards spatial alternation may continue to interfere with odor-cued choices even after the mice performed above chance in the odor-cued version. To test this possibility, we analyzed the four different combinations of trial types – alternating correct, alternating incorrect, non-alternating correct, non-alternating incorrect – with correct and incorrect referring to the odor-guided response and alternating and non-alternating referring to the turn direction compared to the previous choice. Although not rewarded, alternation behavior was above chance (*n* = 8 mice, Z = 2.28, p = 0.011, one-tailed Wilcoxon signed-rank test). However, there was no interaction with the odor-guided responses (correct odor-guided responses in 68.8% of trials; alternation behavior in 63.8% of trials; 𝜒^2^(1, 1207) = 0.01, p = 0.92).

Given that we observed choices that were guided by the odor and also choices that were consistent with alternation above chance, we analyzed the LFP signal that occurred immediately preceding the choice point (i.e., on the stem arm) across different types of trials. Coherence between pairs of regions in the mPFC-dHC-vHC network in the stem arm was not different between trials with correct and incorrect odor-cued responses (*n* = 831 correct and 376 incorrect trials, mPFC-dHC: Z = -0.48, p = 0.63; mPFC-vHC: Z = -0.05, p = 0.96; dHC-vHC: Z = -0.77, p = 0.44, Wilcoxon signed-rank test; **Figure 7A**). However, we found that coherence was significantly higher during trials with alternating choices compared to same-side (i.e., non-alternating) choices irrespective of whether they were correct or incorrect with respect to the odor (*n* = 771 alternating and 436 non-alternating trials, mPFC-dHC: Z = 2.99, p = 0.003; mPFC-vHC: Z = 2.37, p = 0.02; dHC-vHC: Z = 2.28, p = 0.02, Wilcoxon signed-rank test; **Figure 7B**). These results are consistent with reports that prefrontal-hippocampal theta synchrony increases at the choice-point of a Y- or W-maze during a spatial alternation tasks (Benchenane et al., 2010; Jones & Wilson, 2005; Tavares & Tort, 2021). Coherence of canonical theta in the prefrontal-hippocampal network is therefore unrelated to odor-guided choices but rather seems to reflect the extent to which the mice use spatially guided behavior.

**Figure 7.**
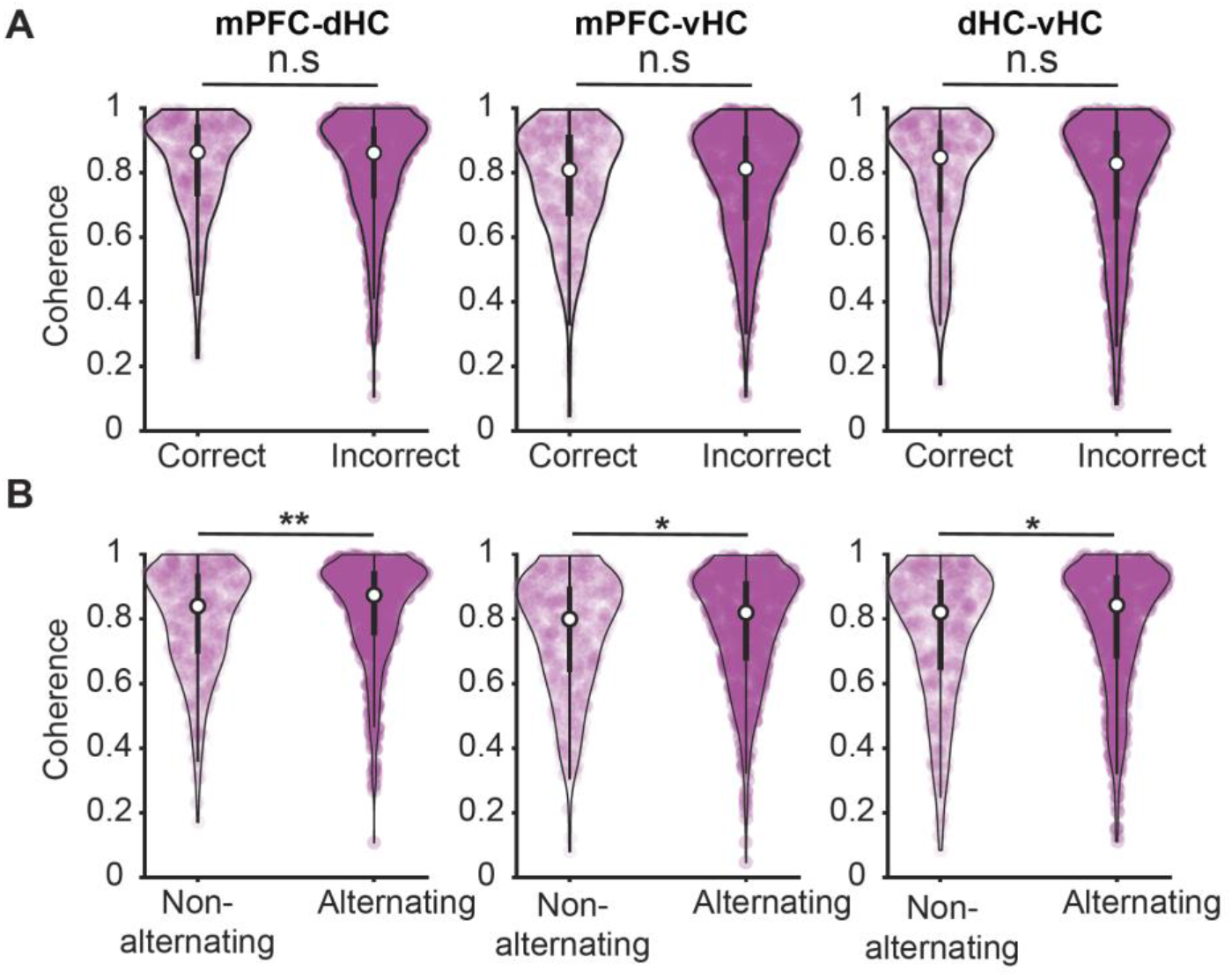
Movement-related theta oscillations in the stem arm were more coherent when mice alternated between left and right arms than when choosing the same arm. **A.** Coherence of movement-related theta oscillations between pairs of regions in the mPFC-dHC-vHC network in the stem arm is compared between trials with correct and incorrect choices. Coherence was not different between trials with correct and incorrect choices (*n* = 831 correct and 376 incorrect trials, mPFC-dHC: Z = -0.48, p = 0.63; mPFC-vHC: Z = -0.05, p = 0.96; dHC-vHC: Z = -0.77, p = 0.44, Wilcoxon signed-rank test). **B.** Same as A, but for alternating compared to non-alternating choices. Coherence was significantly higher during trials with alternating choices compared to trials with non-alternating choices (*n* = 771 alternating and 436 non-alternating trials, mPFC-dHC: Z = 2.99, p = 0.003; mPFC-vHC: Z = 2.37, p = 0.02; dHC-vHC: Z = 2.28, p = 0.02, Wilcoxon signed-rank test). In the violin plots, the center circle indicates the median, and the bottom and top of the thick black lines indicate the 25^th^ and 75^th^ percentile of the data, respectively.

## DISCUSSION

Synchronized oscillations are thought to facilitate coordinated computations across brain regions. Although it is well established that respiration-entrained oscillations are propagated from the OB to other cortical areas, it is unclear to what extent these respiration-entrained oscillations get coupled to endogenous theta oscillations in the prefrontal-hippocampal regions. If coupling occurs, it would be an indication that oscillations that are generated by two different mechanisms in the brain could dynamically couple to support memory computations. To investigate the coordination of respiration-entrained oscillations in the OB and of theta oscillations in the prefrontal-hippocampal circuit, we analyzed simultaneously recorded LFP signals from the OB, mPFC, dHC and vHC during an odor-cued working memory task. We found that respiration-entrained oscillations in the OB were distributed across the 3-12 Hz frequency range within each task phase, including phases when animals moved and phases when animals were predominantly immobile. By examining these task phases separately, we were able to test whether movement-related and sensory-evoked theta oscillations in the prefrontal-hippocampal circuit couple with respiration-entrained oscillations mediated by the olfactory bulb. For movement-related theta, we found that OB and cortical oscillations either occurred at different frequencies, or when occurring at similar frequencies, remained uncoupled from respiration-entrained oscillations in the OB. A similar result was also observed for sensory-evoked theta oscillations during odor sampling periods. Taken together, respiration-entrained oscillations that were propagated from the OB to prefrontal-hippocampal regions did not become coupled to local theta oscillations even during odor sampling periods when olfactory inputs can be assumed to strongly drive information processing and sensory-evoked theta. RROs in the prefrontal-hippocampal regions therefore remained independent from canonical theta oscillations and increased coupling of the oscillations did not appear to serve as a conduit for information processing in an odor-cued working memory task. This was even observed for ventral hippocampal recording locations, which have strong connections with mPFC and piriform cortex and which have been implicated to be an important brain region for odor working memory (Hoover & Vertes, 2007; Kesner et al., 2011). Rather than finding a relation between odor-cued behavior and the coherence between RROs and theta, we found that movement-related theta oscillations were more coherent within the prefrontal-hippocampal network when the spatial component of the task guided the behavioral responses.

### Breathing frequency is variable but only weakly controlled by ongoing behavior

It has long been known that rodent breathing frequencies can vary over a wide range. Mice have a “passive” breathing frequency of 1-4 Hz during quiescence (Jessberger et al., 2016; Wesson et al., 2011). Upon exposure to a novel odor, mice begin “active” sniffing at a high frequency of 4-12 Hz (Jessberger et al., 2016; Wesson et al., 2008; Wesson et al., 2011). Such a modulation of respiration frequencies during odor sampling has been thought to be the basis for odor processing in lower-order olfactory circuits (Wesson et al., 2008). Indeed, sniffing frequency changes the number of odor molecules arriving at the olfactory sensory neurons, thereby increasing their responsiveness to odors at these higher sniffing frequencies (Courtiol et al., 2011). However, further investigations of the role of sniffing frequencies in odor information processing has revealed that mice have varied strategies in terms of sniffing frequencies (Reisert et al., 2020; Wesson et al., 2008). While sniffing frequencies increase in response to a novel odor sampling, mice are able to perform “easy” as well as “difficult” odor discrimination tasks without a significant increase in their sniffing frequencies compared to baseline (Wesson et al., 2008; Wesson et al., 2009). Moreover, the relationship between sniffing frequencies and odor guided behavior is confounded by multiple factors such as locomotion and reward expectation, both of which lead to increased sniffing rates (Bramble & Carrier, 1983; Clarke, 1971; Hérent et al., 2020; Wesson et al., 2008). Our results are consistent with a weak control of breathing frequencies by ongoing behavior because we find that a similarly broad range of OB oscillations can occur in any of the behavioral phases in an odor-cued working memory task.

### Coupling of movement-related theta and sensory-evoked theta to RROs

Although oscillations in the 4-12 Hz band are broadly referred to as theta, it is well established that theta oscillations in the hippocampus are of at least two types – type I and type II. Type I theta is atropine-insensitive and is movement-related (Kramis et al., 1975; Vanderwolf, 1969). Power and frequency of type I theta oscillations have been shown to increase with higher running speeds (Feder & Ranck, 1973; Kuo et al., 2011). Our analysis of movement-related theta oscillations in the return and stem arms revealed that theta oscillations during those periods showed similar relations to movement as type I theta (**Figure 3 – figure supplement 1A**). On the other hand, type II theta is atropine sensitive and is unrelated to movement (Kramis et al., 1975; Vanderwolf, 1969). Type II theta is elicited when the animal is exposed to arousing, vigilant and aversive conditions, such as a predator’s smell (Sainsbury et al., 1987). Our recording of sensory-evoked theta oscillations during the odor sampling period is akin to type II theta oscillations although we did not test the atropine sensitivity of these oscillations. However, we confirmed that these theta oscillations occur while the mouse’s nose was held in the odor port and stationary (**Figure 3A and B and Figure 3 – figure supplement 1B**), which suggests that theta oscillations during this task phase fulfill at least one of the criteria for type II theta. By including task phases in the analysis when theta was either movement-related or sensory-evoked, we were therefore able to test to what extent each type of theta was related to RROs.

For movement-related theta oscillations in the hippocampus, seminal work that recorded nasal air flow as well as LFP from the OB and dorsal hippocampus found that the hippocampal movement-related theta does not couple with the respiration rhythm during exploration (Vanderwolf & Szechtman, 1987). Similarly, RROs were identified as a separate oscillation from theta oscillation during running based on differences in the depth profiles across hippocampal recording sites between both types of oscillations (Nguyen Chi et al., 2016). These reports correspond to our finding that movement-related theta and RROs remain separate oscillations during all phases of behavior. However, when investigating coupling between the two types of oscillations, Tort et al. (2018) reported that theta oscillations within the hippocampus and mPFC were coherent with the respiration rhythm during exploration. While it is therefore possible that RROs become the predominant type of oscillation at a subset of recording sites (e.g., in dentate gyrus), we show that there are also additional prefrontal and hippocampal high-amplitude theta oscillations that do not couple to OB oscillations and are restricted to a more narrow frequency band than RROs.

The evidence for coupling between RROs and sensory-evoked theta during odor sampling is stronger than for movement-related theta but also not equivocal. For example, Macrides et al. (1982) reported that during odor sampling, theta oscillations in the hippocampus couple with the respiration rhythm during the initial stages of learning an odor discrimination reversal task, but that coherence between these oscillations was low in expert animals. Conversely, Kay (2005) showed that hippocampal theta oscillations and the sniffing rhythm were coherent during odor sniffing in a two-odor discrimination task and that the coherence was positively correlated to performance, which suggests that coherence remained high even when animals were proficient. These discrepancies could, at least in part, be explained by the consideration that both canonical theta and RROs can be recorded with hippocampal electrodes. What is interpreted as coherence between hippocampal theta and respiration-entrained OB oscillations could therefore be coherence between OB oscillations and RROs that can be recorded in the hippocampus. Such hippocampal RROs are readily detectable in previous studies and in our data and are particularly pronounced for electrodes in the dentate gyrus (Nguyen Chi et al., 2016; Yanovsky et al., 2014) and at ventral hippocampal sites (see **Figure 4**). If we were to focus on the RRO component of our LFP signals, we would indeed find that it is strongly coupled to OB oscillations during odor sampling (see **Figure 4C**), similar to what has been reported during immobility (Nguyen Chi et al., 2016; Tort et al., 2018). However, we also find that RROs are not the only type of oscillation that can be recorded in hippocampus and mPFC during odor sampling. A second type of oscillation was revealed that showed a narrower frequency distribution than RROs at the same recording sites and corresponded to canonical theta oscillations during the odor sampling phase (see **Figure 3B**).

Our results therefore concur with existing narratives that respiration-entrained oscillations are detected in the prefrontal-hippocampal areas and can be particularly evident when the respiration frequency is lower than the theta oscillation frequency in the hippocampus (Nguyen Chi et al., 2016). In fact, a majority of investigations have studied periods of low respiration frequency (<6 Hz) (Lockmann et al., 2016; Nguyen Chi et al., 2016; Yanovsky et al., 2014) while respiration frequencies can be well above 10 Hz and thus be higher than the canonical theta frequency. However, while mitral and tufted cells in the OB are entrained to the respiration rhythm at low frequencies (up to 6 Hz), they fire tonically at higher respiratory frequencies (6-12 Hz) (Kay & Laurent, 1999). Could it therefore be the case that respiration-entrained oscillations in the OB are transmitted differently to downstream cortical areas based on frequency? In the subset of trials with non-overlapping RRO and theta frequencies when RRO frequency was higher than 6 Hz, respiration-entrained OB oscillations did not show increased coherence with hippocampal oscillations at the RRO frequency, which differs from the pronounced coherence when RRO frequency was lower than 6 Hz. Our results are therefore consistent with those by Kay & Laurent (1999), who suggested that higher OB oscillation frequencies may not be forwarded to cortical areas due to the varied entrainment of OB cells to the respiration rhythm. In addition to a lack of coupling that arises from separate pacemakers – nasal air flow for RROs (Onoda & Mori, 1980; Phillips et al., 2012; Wu et al., 2017) and medial septal area for canonical theta (Bland & Bland, 1986; Gaztelu & Buño, 1982; Mitchell et al., 1982) – coupling could also be lost by the limited propagation of RROs to cortical networks once the respiration rhythm as high in frequency as the predominant hippocampal theta frequency.

### Are coupled oscillators across brain regions related to behavioral performance?

Previous studies firmly established that RROs propagate from the OB to downstream brain regions in a variety of brain states including anesthesia, mobility and immobility (Biskamp et al., 2017; Fontanini et al., 2003; Ito et al., 2014; Lockmann et al., 2016; Nguyen Chi et al., 2016; Yanovsky et al., 2014). Based on these observations, it was speculated that respiration-entrained oscillations are a global signal that synchronizes activity across multiple brain regions and supports sensorimotor integration in a context dependent manner (Lockmann et al., 2016; Macrides et al., 1982; Nguyen Chi et al., 2016; Tort et al., 2018; Yanovsky et al., 2014). However, at least one previous study that tested functional coupling at the theta frequency did not find coherent oscillations between the OB and hippocampus. Because these results were obtained in a simple hippocampus-independent odor discrimination task (Fortin et al., 2002), we considered the possibility that coupling between OB and hippocampal oscillations could emerge in a task that involves the learning of associations between odors and spatial locations, which has been shown to be hippocampus-dependent (Gilbert & Kesner, 2004). However, we found that RROs were not coupled to theta oscillations in either the hippocampus or mPFC during any task phase including during odor sampling when the encoding of sensory information to support working memory was hypothesized to most likely engage hippocampal computations. Given that we did not detect coupling between respiratory-related oscillations and canonical theta oscillations, we conclude that each type of oscillation supports a different set of computations and that coupling of sensory to memory processing occurs in a different frequency band and/or different anatomical pathway, such as by beta oscillations in the lateral entorhinal cortex (Igarashi et al., 2014). Our results therefore imply that oscillatory patterns, even when co-occurring in the same brain regions at overlapping frequencies in a task that engages these brain regions, can continue to support distinct computations.

## MATERIALS AND METHODS

### Subjects

Eight mice (VGAT-cre 129S6(FVB)-Slc32a1^tm2(cre)Lowl^/MwarJ, Jackson Labs; *n* = 4 male, *n* = 4 female) that were 4 months old and weighed 20-30 grams were used as subjects. Sample sizes were determined based on the number of mice used in previous studies with recordings of RROs in awake behaving mice. All mice were single-housed in a reverse 12 hr dark/light cycle (lights off at 8 am). Mice were restricted to 85-90% of their *ad libitum* weight and given full access to water. All the training and testing was conducted during the dark phase. All procedures were conducted in accordance with the University of California, San Diego Institutional Animal Care and Use Committee.

### Surgery

Mice were anesthetized with isoflurane (induction: 3%, maintenance: 1.5-2%) and mounted in a stereotaxic frame (David Kopf Instruments, Model 1900). The scalp was cleaned and retracted using a midline incision and the skull was leveled between bregma and lambda. Five holes were drilled in the skull to attach anchor screws. A hole was drilled above the cerebellum to place the ground screw. Craniotomies were performed over four brain regions on the right hemisphere [OB: +4.2 mm anteroposterior (A/P), 0.6 mm mediolateral (M/L); mPFC: +1.8-2 mm A/P, 0.4 mm M/L; dHC: -1.9 mm A/P, 2.0 mm M/L; vHC: -3.3 mm A/P, 3.5 mm M/L] and dura was removed. Wires were implanted in the four brain regions [OB: -1.2 mm dorsoventral (D/V); mPFC: -1.4 mm D/V; dHC: -1.8 mm D/V; vHC: -3.5 mm D/V] to record local field potentials. The wires were threaded through a circuit board with a connector, and the implant was secured with dental cement. Postoperative care was administered as needed and mice were allowed to recover for a minimum of 5 days.

### Histological procedures

Mice were perfused with 0.1 M phosphate-buffered saline (PBS) followed by 4% paraformaldehyde in PBS solution. Brains were post-fixed for 24 hours in 4% paraformaldehyde and then cryoprotected in 30% sucrose solution for 2 days. Brains were then frozen and sliced into 40 µm coronal sections using a sliding microtome. Sections were mounted on electrostatic slides, stained with cresyl violet and coverslipped with Permount (Fisher Scientific, SP15500) to visualize recording locations. Slides were imaged using a virtual slide microscope (Olympus, VS120).

### Olfactometer and odor delivery

A custom plastic odor port was machined, and two IR LEDs (transmitter and receiver) were placed at the entrance of the odor port to detect nose pokes. These LEDs were connected to an Arduino board (Arduino Mega 2560) which was programmed to detect nose pokes and deliver an odor through a custom-made olfactometer. One of the two odors was pseudo-randomly delivered on each trial, with a minimum interval of 2 s to prevent triggering the odor delivery twice within a single trial. A custom written MATLAB script was used to deliver the odor as well as to send a TTL pulse to the Neuralynx acquisition system to timestamp the odor delivery, nose poke in and nose poke out. A hole was drilled at the bottom of the odor port to deliver the odor at a flow rate of up to 1L/min. Two neutral odors (ethyl acetate and isoamyl acetate) were used in the task. These odors were freshly prepared daily in mineral oil (1:5 ratio by volume).

### Behavior

Mice were trained on an odor-cued working memory task. The room was dimly lit and stable environmental cues were placed. The task was performed on a figure-8 maze that was 50 cm above the ground, 75 cm long, and 50 cm wide with 5 cm wide runways. A custom-made olfactometer was placed on one end of the stem arm. The maze was cleaned with 70% alcohol after each animal used the maze. Animals were trained in phases. On the first day, animals were allowed to freely explore the maze for 10 minutes for habituation. After habituation for one day, animals started the first phase of training. In the first phase, animals were gently guided to the odor port to break an IR beam at the entrance of the odor port upon which an odor was delivered. Animals were required to sniff the odor and run to the other end of the stem arm where they were forced to make the correct choice. They were then rewarded with a single chocolate sprinkle (Betty Crocker Parlor Perfect Chocolate Sprinkles) at the reward zone. Care was taken to place the chocolate sprinkle at the reward location only after the animal made its choice. Animals performed 60 trials per day. Once animals learned to nose poke in the odor port and run to the opposite end of the stem arm without guidance in all 60 trials on two consecutive days, they were ready for the second phase. In the second phase, animals performed the task without guidance to nose poke into the odor port and were given a choice to turn in either direction at the other end of the stem arm. Responses on all 60 trials were recorded and analyzed. The data from the second phase are reported in the Results section.

### Electrophysiological recordings

Local Field Potentials (LFP) were recorded using chronically implanted wires. Implanted wires were connected to a head-mounted preamplifier and via a tether to a 32-channel digital data acquisition system (Neuralynx, Bozeman, MT). Continuous LFP was sampled at 32000 Hz and band-pass filtered between 0.1 and 1000 Hz. Position data of a red and a green LED located on either side of the head-mounted preamplifier were tracked by a video camera at a sampling frequency of 30 Hz to determine the spatial location of the animals while they performed the task.

### LFP Analysis

Raw LFP signals were down-sampled to 2000 Hz and a Morlet wavelet of width ratio = 6 was used to determine the power and phase of the oscillations at 30 log-spaced frequencies in the 3-20 Hz range. An average spectrogram was constructed for each maze zone in each trial. For each frequency, phase differences between pairs of oscillations were calculated, and coherence in each maze zone was computed as the length of the resultant vector of phase differences in that zone. For analysis involving predominant frequencies within a region, a peak was detected in the 3-12 Hz range for RROs and in the 6-12 Hz range for movement-related and sensory-evoked theta oscillations. If no peak was detected, that trial was omitted from analysis for that maze zone.

### Statistics

All statistics were performed using built-in functions in MATLAB (R2019b). Non-parametric tests such as KS and Wilcoxon tests were performed. Corrections for multiple tests were performed using the Holm-Bonferroni method.

### Code availability

Code can be accessed at https://github.com/SunandhaSrikanth.

## ACKNOWLEDGEMENTS

We thank members of the Jill Leutgeb and Stefan Leutgeb labs for valuable comments and suggestions at various stages of this project. We thank Dr Cory Root for help with the custom olfactometer set up. We also thank Mr. Vignesh Srinivasan for help with setting up the odor port. This research was supported by grants from the National Institute of Health (NS102915, NS097772 and MH119179).

## SUPPLEMENTAL FIGURES

**Figure 1 – figure supplement 1.**
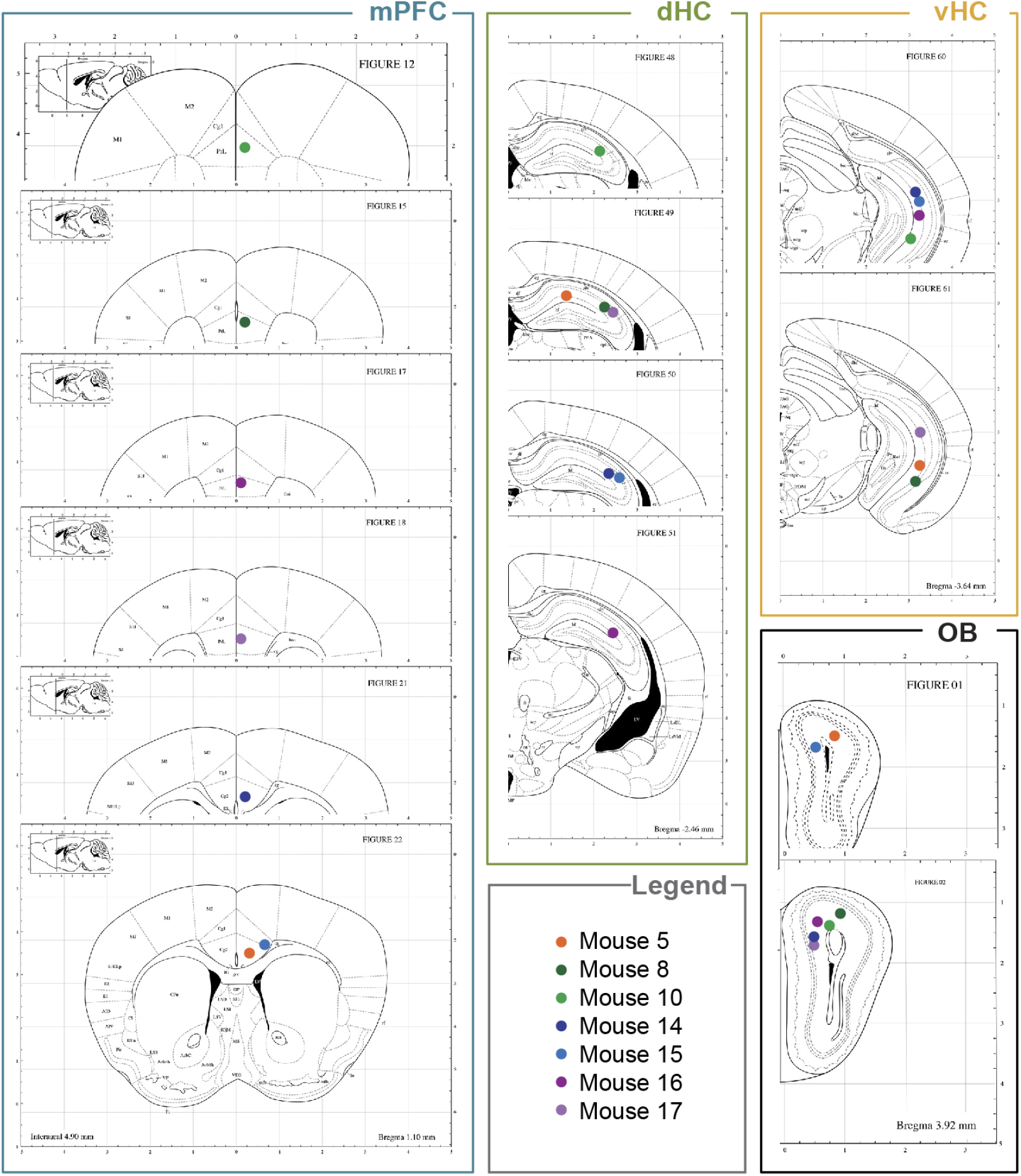
Electrode recording locations. Colored dots indicate the recording locations in each animal (*n* = 7) used in the study. Electrodes were histologically confirmed to be placed in the OB, the prelimbic, infralimbic and anterior cingulate areas of the mPFC and the CA1 areas of the dHC and the vHC.

**Figure 2 – figure supplement 1.**
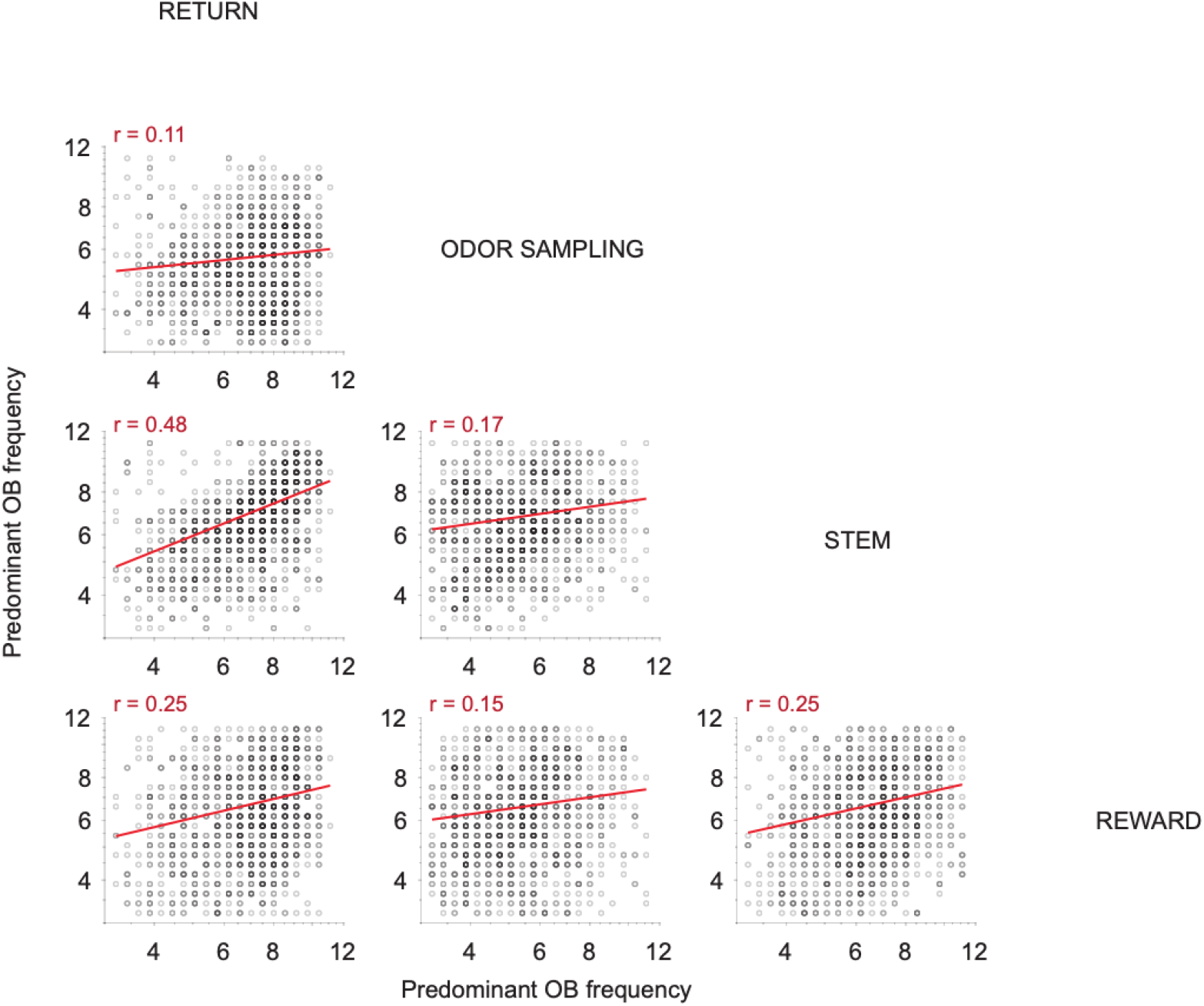
Weak correlations between predominant OB frequencies in different task phases. Scatter plots of predominant OB frequencies during different task phases in each trial. Spearman correlations between predominant OB frequencies during different task phases were weak.

**Figure 3 – figure supplement 1.**
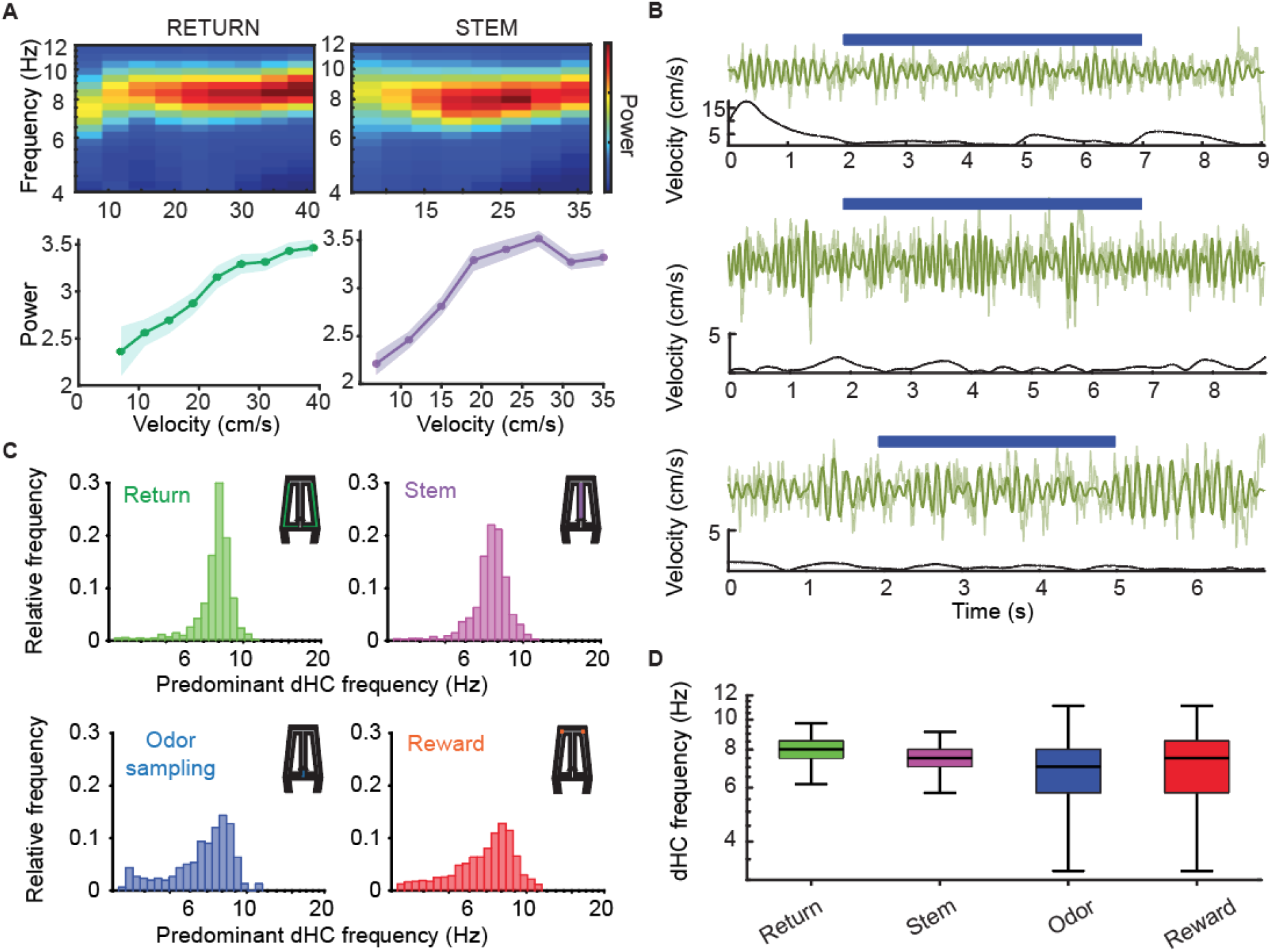
Movement-related and sensory-evoked theta oscillations in the dHC. **A**. (Top) Mean spectrograms during periods of running on the return arms and stem arm, displaying the distribution of power and frequency of movement-related theta oscillations across running speeds. (Bottom) Corresponding average power (±SEM) across running speeds. For each run through the return or the stem arm, the velocity of the animal and dHC power spectrum were calculated and the data was averaged across trials. **B**. Example raw (light green) and 3-12 Hz filtered (dark green) LFP traces of sensory-evoked theta oscillations in the dHC are shown in three different trials. Blue bars represent the time of odor sampling. Velocity traces accompanying each raw LFP trace show the velocity of the animal during the same time period. As expected, velocity was near zero while mice where holding their nose in the odor port. **C**. Distributions of predominant dHC LFP frequencies in each task phase. **D**. Box plots of the predominant dHC frequency in each task phase. Data in C and D corresponds to Figure 3B, but more detailed distributions are shown here. In the box plots, the center line shows the median, and the bottom and top edges of the box represent the 25th and 75th percentiles, respectively. The whiskers indicate the most extreme data points not considered outliers.

**Figure 4 – figure supplement 1.**
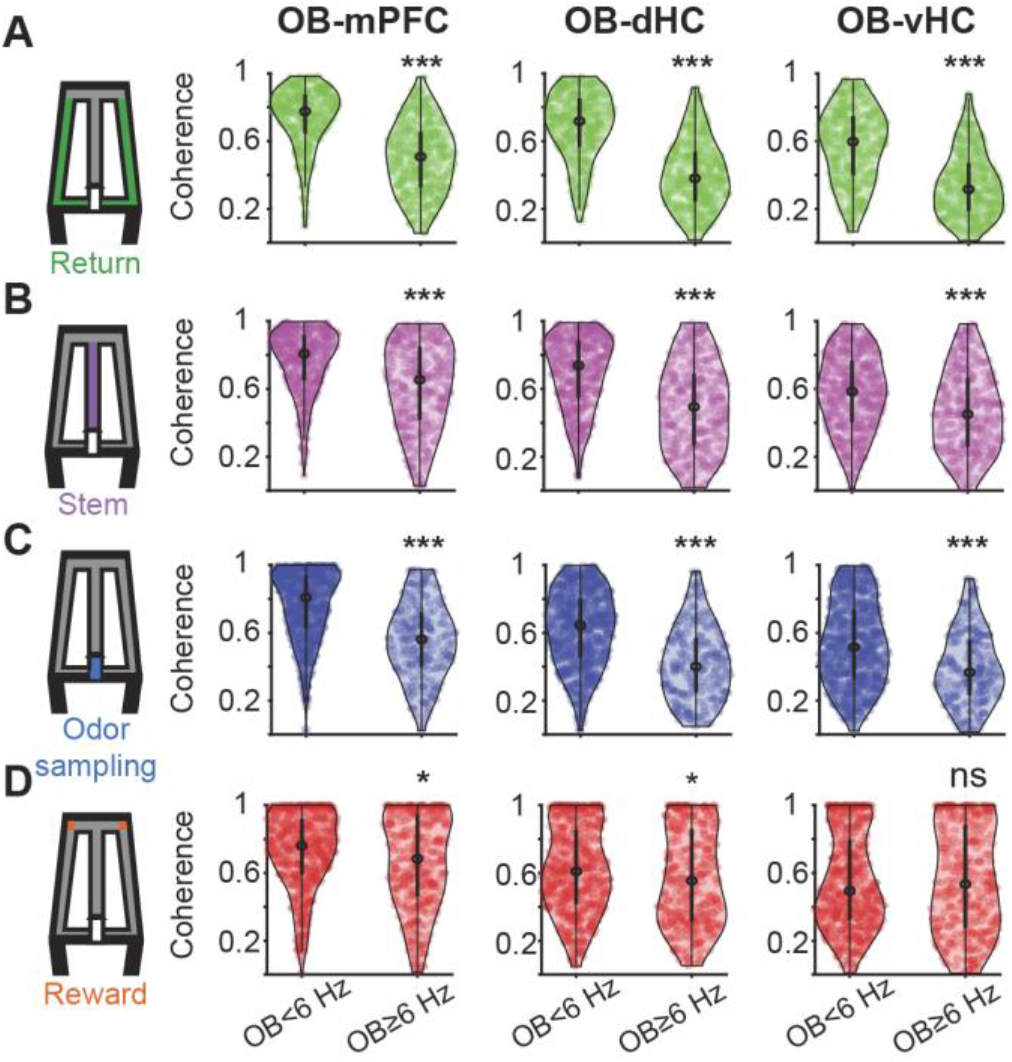
Comparison of coherence between the OB and the prefrontal hippocampal network in non-overlap trials with OB frequency <6 Hz and ≥6Hz. Violin plots of OB-mPFC, OB-dHC, and OB-vHC coherence in (**A**) the return arm, (**B**) the stem arm, (**C**) the odor sampling period and (**D**) the reward arm. Coherence was calculated at the RRO frequency in each trial. Non-overlap trials with OB frequencies <6 Hz and ≥6 Hz were analyzed separately. A lower coherence was found in non-overlap trials with OB frequency ≥6Hz compared to non-overlap trials with OB frequency <6Hz for all combinations of OB recordings with other brain regions in the return arm (*n* = 365 and 378 trials with OB frequency <6 Hz and ≥6 Hz, OB-mPFC: Z = -14.70, p = 6.3e-49; OB-dHC: Z = -15.91, p = 4.9e-57; OB-vHC: Z = -13.16, p = 1.3e-39, Wilcoxon signed-rank test), stem arm (*n* = 447 and 305 trials, OB-mPFC: Z = -8.00, p = 1.2e-15; OB-dHC: Z = -10.31, p = 6.2e-25; OB-vHC: Z = -5.99, p = 2.0e-9, Wilcoxon signed-rank test) and odor sampling period (*n* = 776 and 212 trials, OB-mPFC: Z = -11.57, p = 5.5e-31; OB- dHC: Z = -10.75, p = 5.4e-27; OB-vHC: Z = -7.22, p = 5.1e-13, Wilcoxon signed-rank test). In the reward zone, only OB-mPFC (*n* = 517 and 338 trials, Z = -2.61, p = 0.009, Wilcoxon signed-rank test) and OB-dHC (*n* = 517 and 338 trials, Z = -2.64, p = 0.008, Wilcoxon signed-rank test) coherence was decreased in non-overlap trials with OB frequency ≥6 Hz compared to <6Hz. In the violin plots, the center circle indicates the median, and the bottom and top of the thick black lines indicate the 25^th^ and 75^th^ percentile of the data, respectively.

## REFERENCES

Backus, A., Schoffelen, J., Szebényi, S., Hanslmayr, S., & CF, D. (2016). Hippocampal-Prefrontal Theta Oscillations Support Memory Integration. Current biology : CB, 26(4). https://doi.org/10.1016/j.cub.2015.12.048

Benchenane, K., Peyrache, A., Khamassi, M., Tierney, P. L., Gioanni, Y., Battaglia, F. P., & Wiener, S. I. (2010). Coherent theta oscillations and reorganization of spike timing in the hippocampal-prefrontal network upon learning. Neuron, 66(6), 921–936. https://doi.org/10.1016/j.neuron.2010.05.013

Biskamp, J., Bartos, M., & Sauer, J.-F. (2017). Organization of prefrontal network activity by respiration-related oscillations. Scientific Reports, Published online: 28 March 2017; | doi:10.1038/srep45508. https://doi.org/10.1038/srep45508

Bland, S. K., & Bland, B. H. (1986). Medial septal modulation of hippocampal theta cell discharges. Brain Res, 375(1), 102–116.

Bramble, D., & Carrier, D. (1983). Running and breathing in mammals. Science (New York, N.Y.), 219(4582). https://doi.org/10.1126/science.6849136

Buzsáki, G. (2002). Theta oscillations in the hippocampus. Neuron, 33(3). https://doi.org/10.1016/s0896-6273(02)00586-x

Buzsáki, G., & Draguhn, A. (2004). Neuronal Oscillations in Cortical Networks. https://doi.org/10.1126/science.1099745

Clarke, S. (1971). Sniffing and fixed-ratio behavior for sucrose and brain stimulation reward in the rat. Physiology & behavior, 7(5). https://doi.org/10.1016/0031-9384(71)90133-8

Colgin, L. L. (2011). Oscillations and hippocampal-prefrontal synchrony. Current opinion in neurobiology, 21(3). https://doi.org/10.1016/j.conb.2011.04.006

Courtiol, E., Hegoburu, C., Litaudon, P., Garcia, S., Fourcaud-Trocmé, N., & Buonviso, N. (2011). Individual and synergistic effects of sniffing frequency and flow rate on olfactory bulb activity [research-article]. https://doi.org/10.1152/jn.00672.2011. https://doi.org/10.1152/jn.00672.2011

Feder, R., & Ranck, J. B. (1973). Studies on single neurons in dorsal hippocampal formation and septum in unrestrained rats. II. Hippocampal slow waves and theta cell firing during bar pressing and other behaviors. Exp Neurol, 41(2), 532–555.

Feldman, J. L., Del Negro, C. A., & Gray, P. A. (2013). Understanding the rhythm of breathing: so near, yet so far. Annu Rev Physiol, 75, 423–452. https://doi.org/10.1146/annurev-physiol-040510-130049

Fontanini, A., Spano, P., & Bower, J. M. (2003). Ketamine-xylazine-induced slow (< 1.5 Hz) oscillations in the rat piriform (olfactory) cortex are functionally correlated with respiration. J Neurosci, 23(22), 7993–8001.

Fortin, N. J., Agster, K. L., & Eichenbaum, H. B. (2002). Critical role of the hippocampus in memory for sequences of events [OriginalPaper]. Nature Neuroscience, 5(5), 458–462. https://doi.org/10.1038/nn834

Gaztelu, J. M., & Buño, W. (1982). Septo-hippocampal relationships during EEG theta rhythm. Electroencephalogr Clin Neurophysiol, 54(4), 375–387.

Gilbert, P., & Kesner, R. (2004). Memory for objects and their locations: the role of the hippocampus in retention of object-place associations. Neurobiology of learning and memory, 81(1). https://doi.org/10.1016/s1074-7427(03)00069-8

Hoover, W., & Vertes, R. (2007). Anatomical analysis of afferent projections to the medial prefrontal cortex in the rat. Brain structure & function, 212(2). https://doi.org/10.1007/s00429-007-0150-4

Hyman, J. M., Zilli, E. A., Paley, A. M., & Hasselmo, M. E. (2005). Medial prefrontal cortex cells show dynamic modulation with the hippocampal theta rhythm dependent on behavior. Hippocampus, 15(6), 739–749. https://doi.org/10.1002/hipo.20106

Hérent, C., Diem, S., Fortin, G., & Bouvier, J. (2020). Absent phasing of respiratory and locomotor rhythms in running mice. eLife, 9. https://doi.org/10.7554/eLife.61919

Igarashi, K. M., Lu, L., Colgin, L. L., Moser, M.-B., & Moser, E. I. (2014). Coordination of entorhinal-hippocampal ensemble activity during associative learning. Nature, 510, 143–147. https://doi.org/10.1038/nature13162

Ito, J., Roy, S., Liu, Y., Cao, Y., Fletcher, M., Lu, L., . . . Heck, D.H. (2014). Whisker barrel cortex delta oscillations and gamma power in the awake mouse are linked to respiration. Nat Commun, 5, 3572. https://doi.org/10.1038/ncomms4572

Jessberger, J., Zhong, W., Brankačk, J., & Draguhn, A. (2016). Olfactory Bulb Field Potentials and Respiration in Sleep-Wake States of Mice [Research Article]. Neural Plasticity, 2016. https://doi.org/https://doi.org/10.1155/2016/4570831

Jones, M. W., & Wilson, M. A. (2005). Theta Rhythms Coordinate Hippocampal–Prefrontal Interactions in a Spatial Memory Task. https://doi.org/10.1371/journal.pbio.0030402

Kay, L. M. (2005). Theta oscillations and sensorimotor performance. Proc Natl Acad Sci U S A, 102(10), 3863–3868. https://doi.org/10.1073/pnas.0407920102

Kay, L. M., & Laurent, G. (1999). Odor- and context-dependent modulation of mitral cell activity in behaving rats. Nat Neurosci, 2(11), 1003–1009. https://doi.org/10.1038/14801

Kesner, R. P., Hunsaker, M. R., & Ziegler, W. (2011). The role of the dorsal and ventral hippocampus in olfactory working memory. Neurobiol Learn Mem, 96(2), 361–366. https://doi.org/10.1016/j.nlm.2011.06.011

Kramis, R., Vanderwolf, C. H., & Bland, B. H. (1975). Two types of hippocampal rhythmical slow activity in both the rabbit and the rat: relations to behavior and effects of atropine, diethyl ether, urethane, and pentobarbital. Exp Neurol, *49*(1 Pt 1), 58-85.

Kuo, T. B., Li, J. Y., Chen, C. Y., & Yang, C. C. (2011). Changes in hippocampal θ activity during initiation and maintenance of running in the rat. Neuroscience, 194, 27–35. https://doi.org/10.1016/j.neuroscience.2011.08.007

Lockmann, A. L., Laplagne, D. A., Leão, R. N., & Tort, A. B. (2016). A Respiration-Coupled Rhythm in the Rat Hippocampus Independent of Theta and Slow Oscillations. J Neurosci, 36(19), 5338–5352. https://doi.org/10.1523/JNEUROSCI.3452-15.2016

Macrides, F., Eichenbaum, H. B., & Forbes, W. B. (1982). Temporal relationship between sniffing and the limbic theta rhythm during odor discrimination reversal learning. J Neurosci, 2(12), 1705–1717.

Mitchell, S. J., Rawlins, J. N., Steward, O., & Olton, D. S. (1982). Medial septal area lesions disrupt theta rhythm and cholinergic staining in medial entorhinal cortex and produce impaired radial arm maze behavior in rats. J Neurosci, 2(3), 292–302.

Moberly, A. H., Schreck, M., Bhattarai, J. P., Zweifel, L. S., Luo, W., & Ma, M. (2018). Olfactory inputs modulate respiration-related rhythmic activity in the prefrontal cortex and freezing behavior. Nat Commun, 9(1), 1528. https://doi.org/10.1038/s41467-018-03988-1

Nguyen Chi, V., Müller, C., Wolfenstetter, T., Yanovsky, Y., Draguhn, A., Tort, A. B., & Brankačk, J. (2016). Hippocampal Respiration-Driven Rhythm Distinct from Theta Oscillations in Awake Mice. J Neurosci, 36(1), 162–177. https://doi.org/10.1523/JNEUROSCI.2848-15.2016

Onoda, N., & Mori, K. (1980). Depth distribution of temporal firing patterns in olfactory bulb related to air-intake cycles. J Neurophysiol, 44(1), 29–39. https://doi.org/10.1152/jn.1980.44.1.29

Phillips, M. E., Sachdev, R. N., Willhite, D. C., & Shepherd, G. M. (2012). Respiration drives network activity and modulates synaptic and circuit processing of lateral inhibition in the olfactory bulb. J Neurosci, 32(1), 85–98. https://doi.org/10.1523/JNEUROSCI.4278-11.2012

Reisert, J., Golden, G. J., Dibattista, M., & Gelperin, A. (2020). Dynamics of odor sampling strategies in mice. PLoS One, 15(8), e0237756. https://doi.org/10.1371/journal.pone.0237756

Rojas-Líbano, D., Frederick, D. E., Egaña, J. I., & Kay, L. M. (2014). The olfactory bulb theta rhythm follows all frequencies of diaphragmatic respiration in the freely behaving rat. Front Behav Neurosci, 8, 214. https://doi.org/10.3389/fnbeh.2014.00214

Sainsbury, R. S., Heynen, A., & Montoya, C. P. (1987). Behavioral correlates of hippocampal type 2 theta in the rat. Physiol Behav, 39(4), 513–519.

Siapas, A. G., Lubenov, E. V., & Wilson, M. A. (2005). Prefrontal phase locking to hippocampal theta oscillations. Neuron, 46(1), 141–151. https://doi.org/10.1016/j.neuron.2005.02.028

Tavares, L. C. S., & Tort, A. B. L. (2021). Hippocampal–prefrontal interactions during spatial decision-making - Tavares - - Hippocampus - Wiley Online Library. https://doi.org/10.1002/hipo.23394

Tort, A. B. L., Ponsel, S., Jessberger, J., Yanovsky, Y., Brankačk, J., & Draguhn, A. (2018). Parallel detection of theta and respiration-coupled oscillations throughout the mouse brain. Sci Rep, 8(1), 6432. https://doi.org/10.1038/s41598-018-24629-z

Vanderwolf, C. H. (1969). Hippocampal electrical activity and voluntary movement in the rat. Electroencephalogr Clin Neurophysiol, 26(4), 407–418.

Vanderwolf, C. H., & Szechtman, H. (1987). Electrophysiological correlates of stereotyped sniffing in rats injected with apomorphine. Pharmacol Biochem Behav, 26(2), 299–304. https://doi.org/10.1016/0091-3057(87)90122-5

Wesson, D. W., Donahou, T. N., Johnson, M. O., & Wachowiak, M. (2008). Sniffing behavior of mice during performance in odor-guided tasks. Chem Senses, 33(7), 581–596. https://doi.org/10.1093/chemse/bjn029

Wesson, D. W., Varga-Wesson, A. G., Borkowski, A. H., & Wilson, D. A. (2011). Respiratory and sniffing behaviors throughout adulthood and aging in mice. Behav Brain Res, 223(1), 99–106. https://doi.org/10.1016/j.bbr.2011.04.016

Wesson, D. W., Verhagen, J. V., & Wachowiak, M. (2009). Why Sniff Fast? The Relationship Between Sniff Frequency, Odor Discrimination, and Receptor Neuron Activation in the Rat [research-article]. https://doi.org/10.1152/jn.90981.2008. https://doi.org/10.1152/jn.90981.2008

Wu, R., Liu, Y., Wang, L., Li, B., & Xu, F. (2017). Activity Patterns Elicited by Airflow in the Olfactory Bulb and Their Possible Functions. J Neurosci, 37(44), 10700–10711. https://doi.org/10.1523/JNEUROSCI.2210-17.2017

Yanovsky, Y., Ciatipis, M., Draguhn, A., Tort, A. B., & Brankačk, J. (2014). Slow oscillations in the mouse hippocampus entrained by nasal respiration. J Neurosci, 34(17), 5949–5964. https://doi.org/10.1523/JNEUROSCI.5287-13.2014

Zielinski, M. C., Shin, J. D., & Jadhav, S. P. (2019). Coherent Coding of Spatial Position Mediated by Theta Oscillations in the Hippocampus and Prefrontal Cortex. The Journal of neuroscience : the official journal of the Society for Neuroscience, 39(23). https://doi.org/10.1523/JNEUROSCI.0106-19.2019

